# STracking: a free and open-source python library for particle tracking and analysis

**DOI:** 10.1101/2022.03.10.483766

**Authors:** Sylvain Prigent, Cesar Augusto Valades-Cruz, Ludovic Leconte, Jean Salamero, Charles Kervrann

## Abstract

**Summary:** Analysis of intra and extra cellular dynamic like vesicles transport involves particle tracking algorithms. Designing a particle tracking pipeline is a routine but tedious task. Therefore, particle dynamics analysis is often performed by combining several pieces of software (filtering, detection, tracking…) requiring a lot of manual operations, and therefore leading to poorly reproducible results. A good synergy between particle detector with a tracker is of paramount importance. In addition, a user-friendly interface to control the quality of estimated trajectories is necessary. To address these issues, we developed STracking a python library that allows to combine algorithms into standardized particle tracking pipelines.

**Availability and Implementation:** STracking is available as a python library using “pip install” and the source code is publicly available on GitHub (https://github.com/sylvainprigent/stracking). A graphical interface is available using two napari plugins: `napari-stracking` and `napari-tracks-reader`. These napari plugins can be installed via the napari plugins menu or using “pip install”. The napari plugin source codes are available on GitHub (https://github.com/sylvainprigent/napari-tracks-reader, https://github.com/sylvainprigent/napari-stracking).

## 1 Introduction

The study of cell biology dynamics, such as intracellular membrane transport inward (i.e., endocytosis) and outward (i.e., exocytosis/recycling), has been difficult or at least incomplete until recently, due to the heterogeneity of the motion behavior of the analyzed structures. Different tracking software have been published (e.g., uTrack (Jaqaman et al., 2008), Trackmate (Tinevez et al., 2017), MTT (Sergé et al., 2008), MSSEF-TSAKF (Jaiswal et al., 2015), maptrack (Feng et al., 2011), Cell Tracking Profiler (Mitchell et al., 2020), btrack (Ulicna et al., 2021)) to track individual biomolecules, obtain spatial information and quantify their kinetics. Most of them have focused on the accuracy and reproducibility of the analyses but the user interfaces remain complex or even limited to a two-dimensional representation. The development of a user-friendly graphical user interface (GUI) therefore appears necessary to facilitate the selection of parameters, the analysis and the visualization of 3D+time trajectories estimated from complex 3D videos.

The use of Python, a versatile and free programming language (Fernandez-Gonzalez et al., 2022) is growing rapidly within the bioimaging user community. Python tools for visualization (e.g., napari (Sofroniew et al., 2021), ipyvolume (Breddels et al., 2018), SeeVis (Hattab & Nattkemper, 2019)) and analysis (ZeroCostDL4Mic (von Chamier et al., 2021), BioImageIT (Prigent et al., 2021), Cellpose (Stringer et al., 2021)) are widely applied to microscopy images.

On the other hand, a lot of particle tracking approaches have been developed over the last decades. Interestingly, although a number of studies aimed at comparing particle tracking performance have been published, (Carter et al., 2005; Cheezum et al., 2001; Chenouard et al., 2014; Ruusuvuori et al., 2010; Smal et al., 2010; Smal & Meijering, 2015), none of the tested methods seems to perform in a generic way, regardless of the type of image data. As a consequence, it is critical for users to have the possibility to test different detectors and/or trackers in order to identify the best solution for their application.

Here, we present STracking, an open-source Python library for combining algorithms into standardized particles tracking pipelines for biological microscopy images. STracking is distributed under a GNU General Public License v3.0. STracking combines particle detection, tracking and analysis methods and can be used via a napari plugin. STracking contributes to the recent ecosystem of python-based plugins for bioimage analysis.

## 2 Implementation and application

STracking breaks down a particle tracking pipeline into five components: i) frame-by-frame particle detection; ii) particle linking; iii) analysis of particle properties; iv) design of track features; v) filtering of tracks. Each component is represented as a python object.

Each component can be implemented separately. This modular design makes it easy to update and facilitates interoperability with other plugins/algorithms. Whenever a new detection or tracker algorithm is added, compatibility is guaranteed with the particle tracking pipeline, and it is versioned within the STracking library. Several particle detectors are available in Stracking: Difference of Gaussian (DoG), Determinant of Hessian (DoH), and Laplacian of Gaussian (LoG) from the python library scikit-image (van der Walt et al., 2014). Moreover, STracking includes a tracker (Matov et al., 2011) that estimates the optimal tracks as follows: first, a connection graph is created with all the possible connections. Second, tracks are iteratively extracted from the graph using shortest path and graph pruning.

STracking library uses two data structures: “SParticles” to manage the set of detected particles and “STracks” to manage the collection of trajectories. These data structures contain SciPy(Virtanen et al., 2020) objects to store the particles and the tracks. The particles are represented with a 2D numpy array where each row is dedicated to a specific particle and columns are [T, Z, Y, X] for 3D data and [T, Y, X] for 2D data. The properties of particles are stored in a dictionary. Similarly, tracks are stored in a 2D numpy array where each row is dedicated to specific particle and columns are [trackID, T, Z, Y, X] for 3D data and [trackID, T, Y, X] for 2D data. Tracks features and split/merge events are stored using dictionaries. This data representation is the same as napari (Sofroniew et al., 2021) points and tracks layers, making STracking natively compatible with the napari viewer. We thus implemented a STracking napari plugin suite (`napari-tracks-reader`, `napari-stracking`). It provides a graphical interface to create a STracking pipeline without writing python code. STracking could be used as script in Python or napari plugin. The STracking library can be combined with other Python packages to extend STracking functionalities. The napari plugin allows one to perform a full STracking pipeline, or to load detections or tracks from another software such as StarDist(Schmidt et al., 2018), TrackMate (Tinevez et al., 2017) or UTrack (Jaqaman et al., 2008) and continue the analysis with STracking and napari. Documentation on STracking library with examples, is available at https://sylvainprigent.github.io/stracking/. STracking documentation was created using sphinx and the autodoc extension.

STracking pipeline using the napari plugin is illustrated with data obtained in Lattice Light-Sheet Structured Illumination Microscopy (Chen et al., 2014) (Figure 1 and Supplementary Video 1). The STracking workflow could also be implemented using Jupyter notebook (Supplementary Note 1). Additionally, it can be used for cell migration experiment (Supplementary Note 2). These examples demonstrate the ability of STracking to analyze complex datasets acquired with most advanced microscopy technologies.

**Fig. 1.**
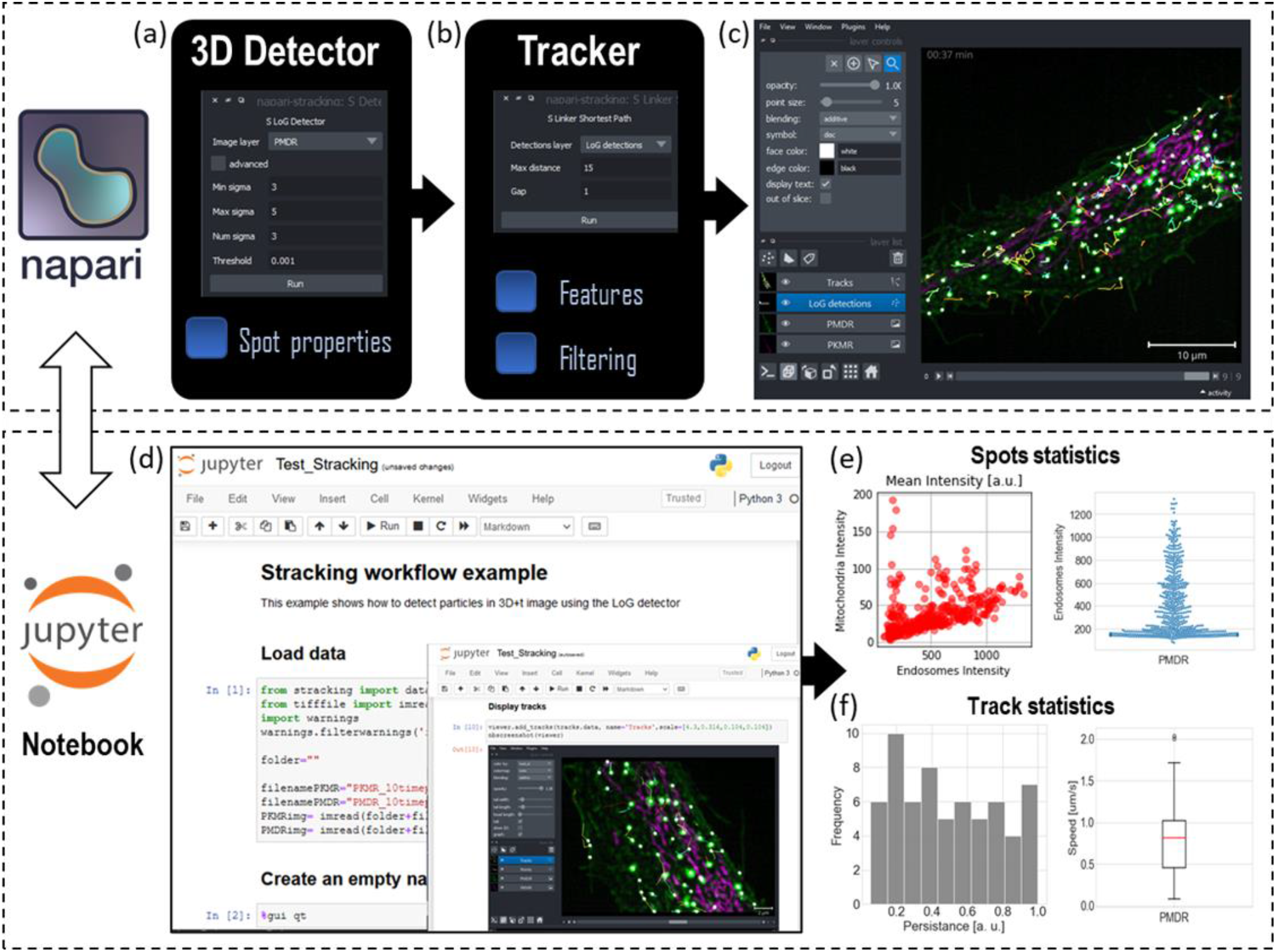
Overview of STracking library implemented in its napari plugin (a-c) and Jupyter Notebook (Kluyver et al., 2016) (d-f). 55 planes 3D volumes of live RPE1 cells double stained with PKMR for Mitochondria (magenta) and with plasma membrane deep red (PMDR) for endosomal pathway(green) were acquired within 4.3s per stack using Lattice light-sheet Structure Illumination Microscopy (LLS-SIM). STracking workflow is illustrated here with single particle tracking of endosomal pathway (PMDR). napari-stracking includes spots detection (a) and linking (b) through a graphical user interface. 3D data and tracks are rendered using napari viewer (c). Jupyter notebook (d) allows also spots (e) and tracks (f) analysis. Additionally, they permit to get spots properties and tracks features, as well as tracks filtering. LLS-SIM data was reconstructed using MAP-SIM (Křížek et al., 2016).

## 3 Conclusions

The STracking library simplifies the design of single particle tracking workflows through a graphical interface using napari and a comprehensive python library of functions. In summary, STracking is a generic single particle tracking library leveraging the 3D visualization capabilities of napari. Unlike previous single particle tracking tools, it provides a very flexible solution for processing and visualizing the tracks that best meet the needs of users. Thus, reproducible analysis can be performed without being an expert programmer. Therefore, our STracking library greatly simplifies the inspection and optimization of single particle tracking algorithms and thus allows the evaluation of new detection and tracker algorithms in this context, which are constantly being developed.

## Data Availability

The data underlying this article are available in FigShare, at https://doi.org/10.6084/m9.figshare.19322171

