## Supplementary Note 1 for "STracking: a free and open-source python library for particle tracking and analysis"

| Supplementary Data | Title |
|---|---|
| **Supplementary Note 1** | Jupyter Notebook using the STracking library in LLS-SIM data. |
| **Supplementary Note 2** | Jupyter Notebook using StarDist(Detection) and the STracking library (tracker) for tracking of cell nuclei. |
| **Supplementary Video 1** | Video example using GUI interface. |

**Supplementary Note 1:** Jupyter Notebook[1] using the STracking library in LLS-SIM data. This notebook corresponds to the workflow of Fig. 1 d-f. Analysis used the libraries matplotlib[2] and seaborn[3].

### Stracking workflow example

This example shows how to detect particles in 3D+t image using the LoG detector

#### Load data

In [1]:
```
from stracking import data
from tifffile import imread

folder=""

filenamePKMR="PKMR_10timepoints_crop.tif"
filenamePMDR="PMDR_10timepoints_crop.tif"
PKMRimg= imread(folder+filenamePKMR)
PMDRimg= imread(folder+filenamePMDR)
```

#### Create an empty napari viewer

In [2]:
```
%gui qt
```

In [3]:
```
import napari
from napari.utils import nbscreenshot
viewer = napari.Viewer(axis_labels='tzyx')
```

#### Display volumetric timeseries

In [4]:
```
viewer.add_image(PKMRimg, name='PKMR', multiscale=False, scale=[4.3,0.316,0.104,0.104],
            contrast_limits=[10, 600], colormap='magenta',blending='additive');

viewer.add_image(PMDRimg, name='PMDR', multiscale=False, scale=[4.3,0.316,0.104,0.104],
            contrast_limits=[10, 1_000], colormap='green',blending='additive',gamma=0.8);

viewer.dims.ndisplay = 3
viewer.scale_bar.visible='true'
viewer.scale_bar.unit='um'
nbscreenshot(viewer)
```

Out[4]:

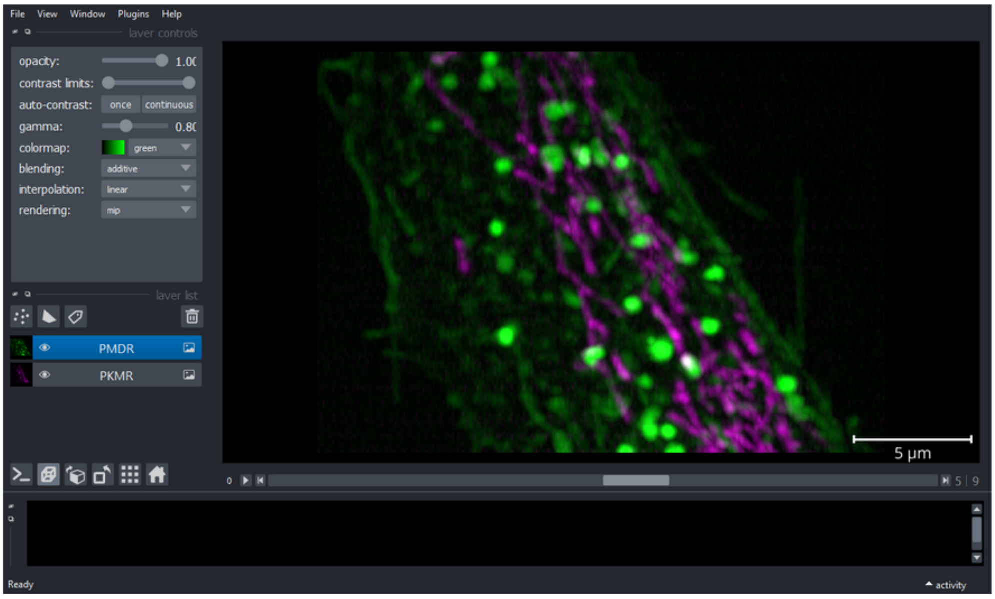

#### LoG 3D+t detection

In [5]:
```
from stracking.detectors import LoGDetector

detector = LoGDetector(min_sigma=3, max_sigma=5, num_sigma=3, threshold=0.001)
particles = detector.run(PMDRimg)
```

#### Display spots

In [6]:
```
viewer.add_points(particles.data, size=5, shading='spherical',scale=[4.3,0.316,0.104,0.104],blending='additive')
nbscreenshot(viewer)
```

Out[6]:

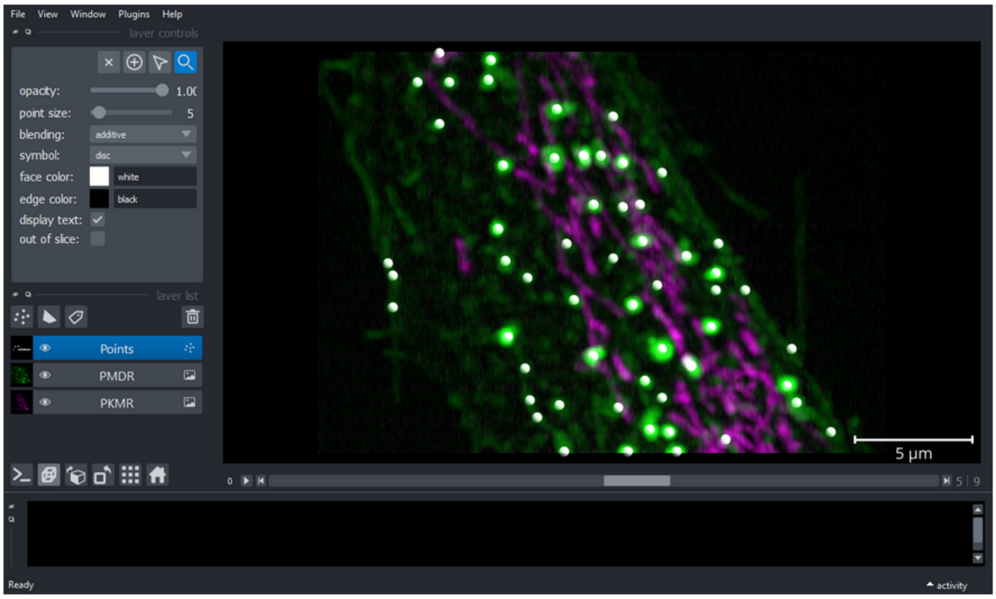

### Spots properties

In [7]:
```
from stracking.properties import IntensityProperty

property_calc = IntensityProperty(radius=2.5)
property_calc.run(particles,PKMRimg)
y=particles.properties['mean_intensity']
particlesch1=particles
property_calc = IntensityProperty(radius=2.5)
property_calc.run(particles,PMDRimg)
x=particles.properties['mean_intensity']
```

#### Spots statistics

In [8]:
```
import matplotlib.pyplot as plt
import numpy as np
from matplotlib import colors
from matplotlib.ticker import PercentFormatter
import seaborn as sns

plt.style.use('_mpl-gallery')
fig, axs = plt.subplots(1, 2, figsize=(15, 4), sharey=False)
axs[0].set_title("Mean Intensity [a.u.]",fontsize=14)
axs[0].set_ylabel("Mitochondria Intensity",fontsize=14)
axs[0].set_xlabel("Endosomes Intensity",fontsize=14)
axs[0].scatter(x,y,alpha=0.5,color='r');
axs[0].tick_params(axis='both', labelsize=14)
sns.set_style("whitegrid")
axs[1] = sns.swarmplot(y=x,alpha=0.9,size=3)
axs[1].set_ylabel("Endosomes Intensity",fontsize=14)
axs[1].set_xlabel("PMDR",fontsize=14)
axs[1].set_title("Endosomes Intensity [a.u.]",fontsize=14)
axs[1].tick_params(axis='both',labelsize=14)
```

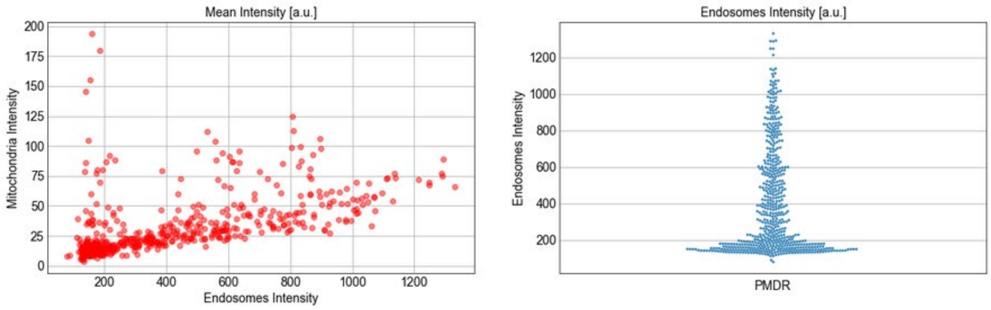

### Tracker

Shortest path tracking with euclidean cost

In [9]:

```
from stracking.linkers import SPLinker, EuclideanCost

euclidean_cost = EuclideanCost(max_cost=225)
my_tracker = SPLinker(cost=euclidean_cost, gap=1)
tracks = my_tracker.run(particles)
```

...

### Display tracks

In [10]:

```
viewer.add_tracks(tracks.data, name='Tracks',scale=[4.3,0.316,0.104,0.104])
nbscreenshot(viewer)
```

Out[10]:

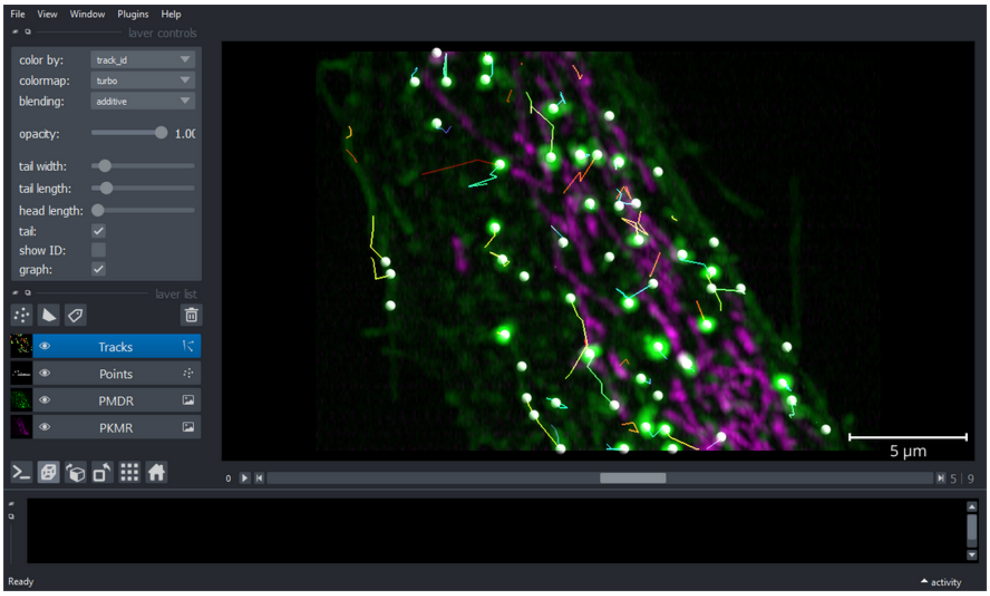

### Features - tracking

In [11]:

```
from stracking.features import (LengthFeature, DistanceFeature,
                                DisplacementFeature)

tracks.scale=[1,0.316,0.104,0.104];

# Length feature
feature_calc = LengthFeature()
tracks = feature_calc.run(tracks)

# Distance feature
feature_calc = DistanceFeature()
tracks = feature_calc.run(tracks)

# Displacement feature
feature_calc = DisplacementFeature()
tracks = feature_calc.run(tracks)
```

#### Display tracks properties

In [12]:
```
displacement=tracks.features['displacement'];
distance=tracks.features['distance'];
length=tracks.features['length'];

displacementMat = np.array([displacement[i] for i in range(len(displacement))])
distanceMat = np.array([distance[i] for i in range(len(distance))])
lengthMat = np.array([length[i] for i in range(len(length))])
speed=distanceMat/(4.3*lengthMat)

fig, axs = plt.subplots(1, 3, figsize=(15, 4), sharey=False)

axs[0].hist(4.3*lengthMat, bins=8, edgecolor="white")
axs[0].set_xlabel("Track Duration [s]",fontsize=14)
axs[0].set_ylabel("Frequency",fontsize=14)
axs[0].tick_params(axis='both', labelsize=14)

axs[1].boxplot([speed],patch_artist=True,
               medianprops={"color": "red", "linewidth": 1.5},
               boxprops={"facecolor": "white", "edgecolor": "black",
                          "linewidth": 1.5},
                whiskerprops={"color": "black", "linewidth": 1.5},
                capprops={"color": "black", "linewidth": 1.5},labels=[''])
axs[1].set_ylabel("Speed [um/s]",fontsize=14)
axs[1].set_xlabel("PMDR",fontsize=14)
axs[1].tick_params(axis='both', labelsize=14)

axs[2].hist(displacementMat/distanceMat, bins=10, edgecolor="white",facecolor='gray',alpha=0.9)
axs[2].set_xlabel("Persistance [a. u.]",fontsize=14)
axs[2].set_ylabel("Frequency",fontsize=14)
axs[2].tick_params(axis='both', labelsize=14)
```

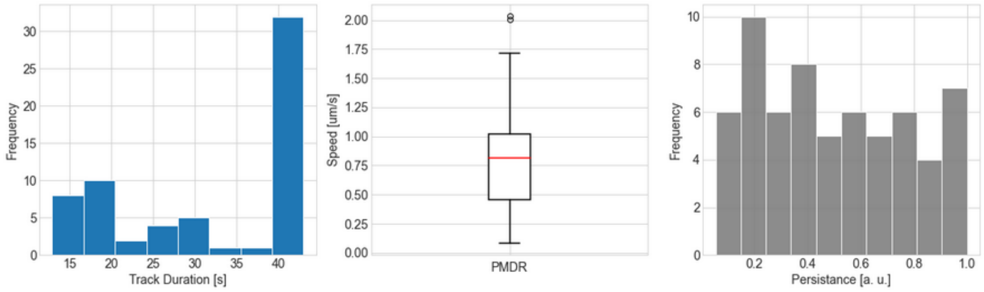

### Features tracking

In [13]:
```
from stracking.filters import FeatureFilter

f_filter = FeatureFilter(feature_name='length', min_val=10, max_val=100)
filtered_tracks = f_filter.run(tracks)

viewer.add_tracks(filtered_tracks.data,features=filtered_tracks.features,name='Filtered Tracks',scale=[4.3,0.316
viewer.layers['Tracks'].visible=False
nbscreenshot(viewer)
```

Out[13]:

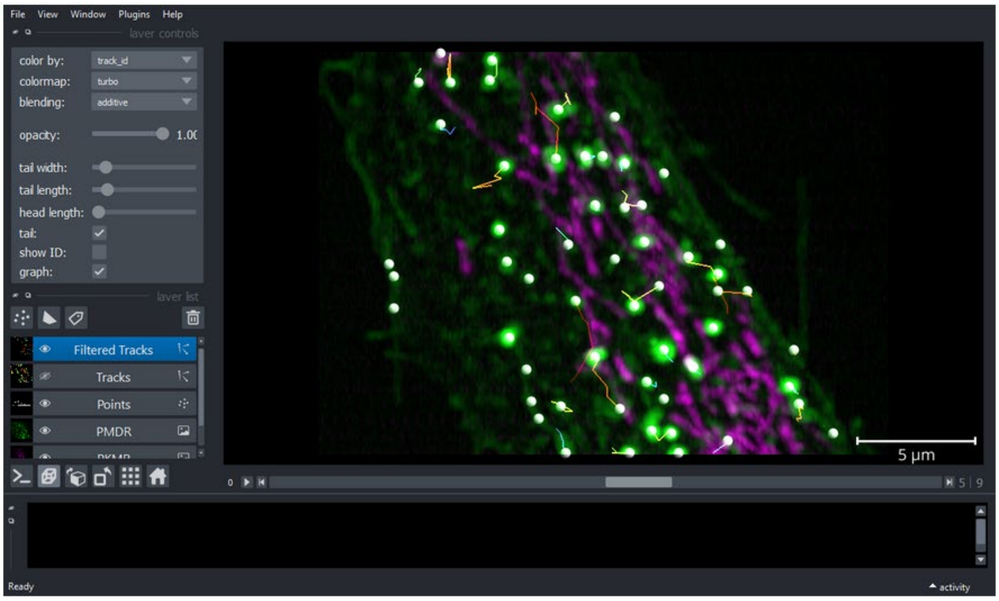

**Supplementary Note 2:** Jupyter Notebook[1] using StarDist[4](Detection) and the STracking library (tracker) for tracking of cell nuclei. Data was extracted from a zenodo repository[5].

### StarDist (Detection) + Stracking (Tracker)

This example shows a combination of StarDist Detection and Stracking (Tracker)

#### Load trained mode

In [1]:
```
from stardist.models import StarDist2D

# prints a list of available models
StarDist2D.from_pretrained()

# creates a pretrained model
model = StarDist2D.from_pretrained('2D_versatile_fluo')
```

```
There are 4 registered models for 'StarDist2D':

Name                   Alias(es)
────                   ────────
'2D_versatile_fluo'    'Versatile (fluorescent nuclei)'
'2D_versatile_he'      'Versatile (H&E nuclei)'
'2D_paper_dsb2018'     'DSB 2018 (from StarDist 2D paper)'
'2D_demo'              None
Found model '2D_versatile_fluo' for 'StarDist2D'.
Loading network weights from 'weights_best.h5'.
Loading thresholds from 'thresholds.json'.
Using default values: prob_thresh=0.479071, nms_thresh=0.3.
```

#### StarDist: Prediction and detection

In [2]:
```
from csbdeep.utils import normalize
import matplotlib.pyplot as plt
from tifffile import imread
import numpy as np

folder=""
filename="P31-crop2.tif"

img= imread(folder+filename)

labels=np.zeros(img.shape);

img=normalize(img)

for i in range(img.shape[0]):
    labels[i,:,:],details = model.predict_instances(img[i,:,:])
    pointstemp= details['points']
    X0 = i*np.ones((pointstemp.shape[0],1))
    pointstemp = np.hstack((X0,pointstemp))
    if i>0:
        points=np.concatenate((points,pointstemp),axis=0)
    else:
        points=pointstemp
```

#### Create an empty napari viewer

In [3]:
```
%gui qt
```

In [4]:
```
import napari
from napari.utils import nbscreenshot
viewer = napari.Viewer(axis_labels='tyx')
```

### Display Input and StarDist Prediction

In [5]:

```
viewer.add_image(img, name='Input', multiscale=False,
                 contrast_limits=[0, 3], colormap='gray',blending='additive');

viewer.add_image(labels, name='Predictions StarDist', multiscale=False,
                 colormap='gist_earth',blending='additive',opacity=0.2);

nbscreenshot(viewer)
```

Out[5]:

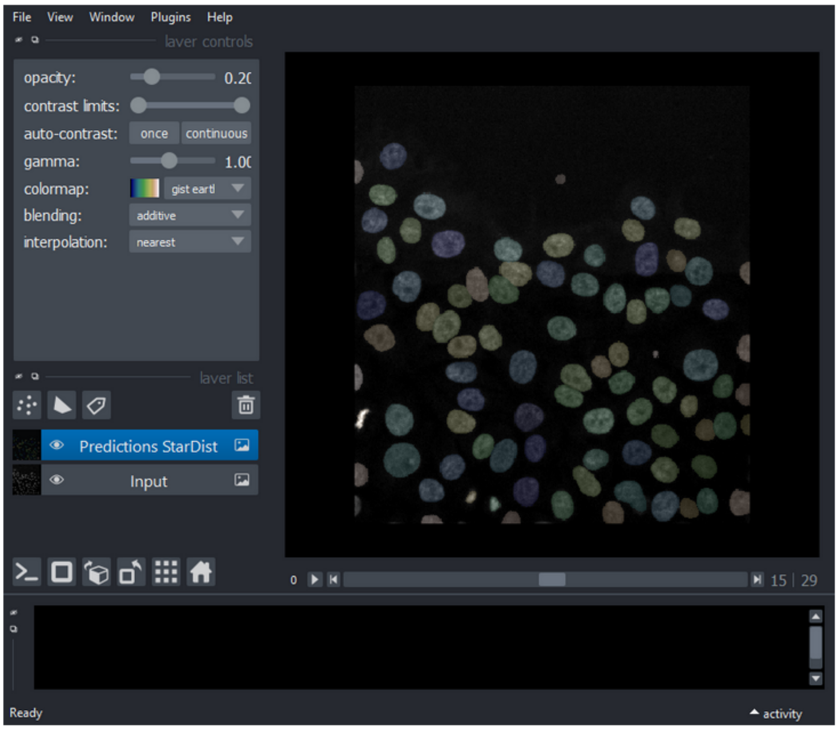

### Display spots from StarDist

In [6]:

```
from stracking.containers import SParticles

particles = SParticles(data=points)
viewer.add_points(particles.data, size=5, blending='additive')
nbscreenshot(viewer)
```

Out[6]:

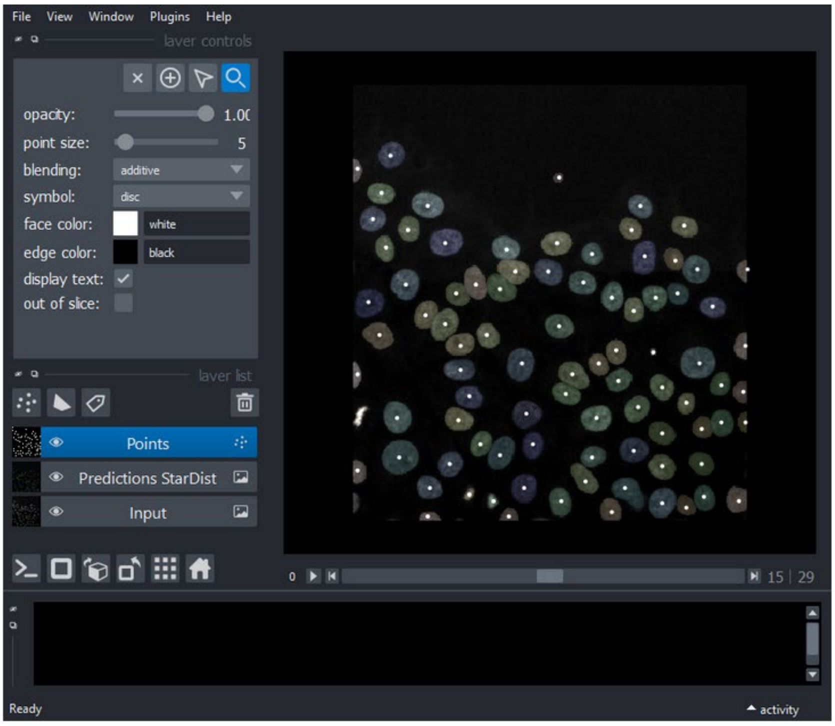

#### Linker

Shortest path tracking with euclidean cost

In [7]:
```
from stracking.linkers import SPLinker, EuclideanCost

euclidean_cost = EuclideanCost(max_cost=100);
my_tracker = SPLinker(cost=euclidean_cost, gap=1);
tracks = my_tracker.run(particles);
```

```
dim in track to path= 2
add predecessor...
add predecessor...
add predecessor...
add predecessor...
extract track...
dim in track to path= 2
add predecessor...
add predecessor...
add predecessor...
extract track...
dim in track to path= 2
add predecessor...
add predecessor...
add predecessor...
extract track...
dim in track to path= 2
add predecessor...
add predecessor...
```

#### Display tracks

In [8]:
```
viewer.add_tracks(tracks.data, name='Tracks')
nbscreenshot(viewer)
```

Out[8]:

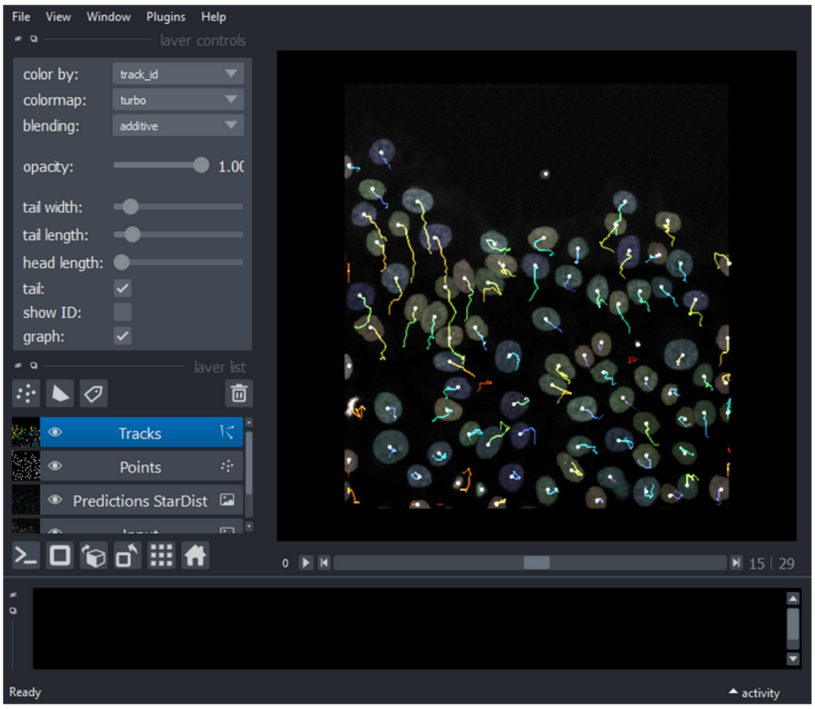
